# Identifying leaf anatomy and metabolic regulators that underpin C_4_ photosynthesis in *Alloteropsis semialata*

**DOI:** 10.1101/2024.03.18.585502

**Authors:** Ahmed S Alenazi, Lara Pereira, Pascal-Antoine Christin, Colin P Osborne, Luke T Dunning

**Author notes:** Corresponding author: Luke T. Dunning; Ecology and Evolutionary Biology, School of Biosciences, University of Sheffield, Western Bank, Sheffield S10 2TN, United Kingdom. These authors contributed equally to the work (Ahmed S Alenazi, Lara Pereira).

## Abstract

- C_4_ photosynthesis is a complex trait requiring multiple developmental and metabolic alterations. Despite this complexity, it has independently evolved over 60 times. However, our understanding of the transition to C_4_ is complicated by the fact that variation in photosynthetic type is usually segregated between species.
- Here, we perform a genome wide association study (GWAS) using the grass *Alloteropsis semialata*, the only known species to have C_3_, intermediate, and C_4_ accessions. We aimed to identify genomic regions associated with the strength of the C_4_ cycle (measured using δ^13^C), and the development of C_4_ leaf anatomy.
- Genomic regions correlated with δ^13^C include regulators of C_4_ decarboxylation enzymes (*RIPK*), non-photochemical quenching (*SOQ1*), and the development of Kranz anatomy (*SCARECROW-LIKE*). Regions associated with the development of C_4_ leaf anatomy in the intermediate accessions contain additional leaf anatomy regulators, including those responsible for vein patterning (*GSL8*) and meristem determinacy (*GRF1*).
- The detection of highly correlated genomic regions with a modest sample size indicates that the emergence of C_4_ photosynthesis in *A. semialata* required a few loci of large effect. The candidate genes could prove to be relevant for engineering C_4_ leaf anatomy in C_3_ species.

## Introduction

Oxygenic photosynthesis originated over two billion years ago and is the ultimate source of nearly all energy used by living organisms. Almost 90% of plants fix carbon using the ancestral C_3_ cycle, but this process is inefficient in hot environments (Sage and Monson, 1999). This is because the key enzyme responsible for the initial fixation of atmospheric CO_2_ (Rubisco) is less able to discriminate CO_2_ from O_2_ at higher temperatures, and as a result energy is lost through photorespiration (Farquhar et al., 1982). To reduce photorespiration plants have evolved C_4_ photosynthesis, replacing the enzyme responsible for initially fixing atmospheric CO_2_ (Edwards & Ku, 1987; Hatch, 1971). In C_4_ species, atmospheric CO_2_ is converted into HCO_3_ by carbonic anhydrase (CA) and then fixed by phosphoenolpyruvate carboxylase (PEPC) into a C_4_ acid. This C_4_ acid is subsequently shuttled into an internal leaf compartment (usually the bundle sheath cells) where it is decarboxylated. The emitted CO_2_ is then re-fixed by Rubisco, which is restricted to this compartment and isolated from atmospheric O_2_. This compartmentalisation effectively prevents photorespiration. C_4_ photosynthesis is a complex trait that relies on both changes to the leaf anatomy and the coordinated regulation of multiple metabolic enzymes (Hatch, 1987). Despite this complexity, C_4_ photosynthesis is a textbook example of convergent evolution, having arisen over 60 times in plants (Sage et al., 2011).

By comparing species with different photosynthetic types, the core C_4_ enzymes, multiple accessory genes, and loci associated with C_4_ leaf anatomy (often termed ‘Kranz’ anatomy) have been identified (Langdale et al., 1987; 1988; Slewinski et al., 2012; Cui et al., 2014). However, decomposing the individual steps during the transition to C_4_ is confounded by the fact that variation in photosynthetic type is usually segregated between distinct species that have been independently evolving for millions of years, meaning that they differ in many aspects besides those linked to the photosynthetic pathway (Heyduk et al. 2019). The interspecific segregation of variation in photosynthetic type makes it challenging to apply quantitative genetics methods, such as quantitative trait loci (QTL) mapping and genome-wide association studies (GWAS), since these rely on traits varying within a species, or the ability to hybridise species with divergent phenotypes. GWAS has been used to investigate the variation of C_4_ traits within C_4_ species, such as photosynthetic performance during chilling in maize (Strigens et al., 2013), and to identify genes associated with stomatal conductance and water use efficiency in sorghum (Ferguson et al., 2021; Pignon et al., 2021). However, to date there has been no QTL region identified for differences in C_4_ carbon fixation or Kranz anatomy (Simpson et al., 2021).

The proportion of carbon that is fixed through the C_4_ cycle is usually measured using the stable carbon isotope ratio (δ^13^C). Both ^12^C and ^13^C occur naturally in the atmosphere, and, in C_3_ plants, Rubisco preferentially fixes ^12^C during photosynthesis. Conversely, in C_4_ plants, carbon is initially fixed by CA and PEPC, and this coupled enzyme system discriminates less than Rubisco between the two isotopes. The rate of CO_2_ release in the bundle sheath is coordinated with the rate of CO_2_ fixation by Rubisco, which reduces the fractionation effect of this enzyme. δ^13^C is therefore commonly used as a proxy for photosynthetic type and the relative strength of the C_4_ cycle. Whilst there is intraspecific variation in δ^13^C for C_4_ species such as maize and *Gynandropsis* (Voznesenskaya et al., 2007), we do not know whether this variation arises from differences in anatomy or biochemistry (Simpson et al., 2022). In addition, some of the observed variation in δ^13^C could also be due to environmental effects on water-use efficiency (Farquhar and Richards, 1984), particularly if the phenotypic data comes from individuals sampled in the field. However, differences in the δ^13^C between accessions of some species are maintained in a common environment (Lundgren et al., 2016), indicating that the δ^13^C ratio likely has a genetic component. Intraspecific, heritable variation in δ^13^C offers an excellent opportunity for using quantitative genetic approaches to discover C_4_ QTLs.

The grass *Alloteropsis semialata* has long been used as a model to study C_4_ evolution, since it is the only species known to have C_3_ and C_4_ genotypes (reviewed by Pereira et al., 2023). This species also has a number of intermediate populations found in the grassy ground layer of the Central Zambezian miombo forests that we refer to as “C_3_+C_4_” because they perform a weak C_4_ cycle in addition to directly fixing CO_2_ through the C_3_ cycle (Lundgren et al., 2016; Dunning et al. 2017). Comparative studies have shown that the transition to a purely C_4_ physiology in *A. semialata* is caused by the overexpression of relatively few core C_4_ enzymes (Dunning et al. 2019a) and the acquisition of C_4_-like morphological traits, notably the presence of minor veins (Lundgren et al. 2019). The δ^13^C of the C_3_+C_4_ plants range from values characteristic of a weak (or absent) C_4_ cycle to values that show that the C_4_ cycle accounts for more than half of the carbon acquisition (von Caemmerer 1992, 2000; Lundgren et al., 2015; Stata and Sage 2019; Olofsson et al. 2021). Furthermore, the strengthening of the C_4_ cycle in the C_3_+C_4_ intermediates is associated with alterations in a number of leaf anatomical traits related to the preponderance of inner bundle sheath tissue, the cellular location of the C_4_ cycle in this species (Alenazi et al., 2023), including the distance between consecutive bundle sheaths, the width of inner bundle sheath cells and the proportion of bundle sheath tissue in the leaf (Alenazi et al., 2023).

*Alloteropsis semialata* therefore represents an ideal system to identify the genetic basis underpinning C_4_ photosynthesis. Here, we first conducted a global analysis to identify candidate genes associated with the strength of the C_4_ cycle (δ^13^C) using genomic data from 420 individuals representing C_3_, C_3_+C_4_ and C_4_ phenotypes. We then focused specifically on the C_3_+C_4_ intermediates, to identify candidate genes associated with the relative expansion of bundle sheath tissue during the transition from a weak to a strong C_4_ cycle. The high level of interspecific variation in *A. semialata* permits a fine-scale understanding of the genetic basis of C_4_ evolution, including the intermediate steps involved in the assembly of this complex trait. This is crucially important to identify the initial changes required for the emergence of this trait, something that may ultimately have applications in the engineering of C_4_ photosynthesis in C_3_ crops such as rice.

## Materials and Methods

### Genome data and population genetic analyses

For the genomic analyses, we compiled previously published double digest restriction-site associated DNA sequencing (ddRADSeq) data sets for *Alloteropsis semialata* (R. Br.) Hitchc. accessions that also had known δ^13^C values (Lundgren et al., 2015 & 2016; Bianconi et al., 2020, Olofsson et al., 2021; Alenazi et al., 2023). In total, the data set comprised 420 individuals from 87 populations across Africa and Asia (Table S1), representing the full range of photosynthetic types found in *A. semialata* (45 x C_3_; 132 x C_3_+C_4_; 243 x C_4_).

The ddRADseq data were downloaded from NCBI Sequence Read Archive and cleaned using Trimmomatic v.0.38 (Bolger et al., 2014) to remove adapter contamination (ILLUMINACLIP option in palindrome mode) and low-quality bases (Q < 3 from both 5ʹ and 3ʹ ends; Q < 15 for all bases in four-base sliding window). The cleaned ddRADseq data were then mapped to a chromosomal scale *A. semialata* reference genome previously assembled for a C_4_ Australian accession (Dunning et al. 2019b) using bowtie2 v.2.2.3 with default parameters (Langmead and Salzberg, 2012). We called SNPs from these alignments using the GATK v3.8 (McKenna et al., 2010; Van der Auwera et al., 2013) pipeline with default parameters. We generated individual variant files (gVCF) with HaplotypeCaller, and then combined them into a single multi-sample VCF file with Genotype GVCFs. Biallelic SNPs were extracted from this file using SelectVariants, and high-quality SNPs retained using VariantFiltration (MQ > 40, QD > 5, FS < 60, MQRankSum > -12.5 ReadPosRankSum > -8). Finally, we used VCFtools to filter remaining SNPs to remove those with > 30% missing data and/or a minor allele frequency < 0.05 (Danecek et al., 2011).

The evolutionary relationship among samples was inferred using a maximum likelihood phylogenetic tree. We used VCF2phylip v.2.8 (Ortiz 2019) to generate a nucleotide alignment from the filtered VCF file. To reduce the effect of linked SNPs on phylogenetic reconstruction, we thinned the data set so that SNPs were at least 1 kb apart (starting from the first SNP on each chromosome). The phylogenetic tree was inferred using RAxML v.8.2.12 (Stamatakis 2014) with the GTRCAT model and 100 bootstrap replicates. Finally, to verify previous phylogenetic groupings (Alenazi et al., 2023), we determined the population structure of the C_3_+C_4_ accessions using Admixture v.1.3.0 (Alexander et al., 2009). We ran the analysis with multiple values of k (range 2 - 7), with 10 replicate runs for each value. The optimal k was inferred using Admixture’s cross-validation error method. We also used PLINK-v1.9 to perform a principal component analysis (PCA) to quantify population structure and to generate a pairwise kinship matrix (Purcell et al. 2007).

### Leaf anatomical traits of C_3_+C_4_ A. semialata

Leaf anatomy data for all 132 C_3_+C_4_ individuals were either extracted from a previous study (n =100; Alenazi et al., 2023) or generated here using the same method (n= 32; Table S2). The measurements themselves were taken from leaf cross-sections that were prepared following the method described by Alenazi et al. (2023). In brief, silica dried leaf material was first rehydrated at 4 °C in 1% KOH solution before being embedded in Technovit 7100 (Heraeus Kulzer GmbH, Wehrhein, Germany). After embedding, 11-μm-thick transverse sections were generated with a rotary microtome (Leica Biosystems, Newcastle, UK), and they were stained for 1.5 minutes with 1% toluidine blue O. (Sigma-Aldrich, St. Louis, MO, USA). The slide images were captured using a mounted camera on an Olympus BX51 microscope (Olympus, Hamburg, Germany), and images from the same leaf were stitched together with Hugin’s software (Hugin Development Team, 2015). All measurements of leaf anatomical characteristics were made using ImageJ v1.53f (Schneider et al. 2012), avoiding the midrib and leaf margins.

We recorded the total cross-sectional areas between secondary veins (i.e. veins accompanied by extraxylary fibers and epidermal thinning) for mesophyll (including airspaces; MS) and inner bundle sheath (IBS) tissues (Figure 1). We used these values to then calculate the inner bundle sheath fraction (IBSF = IBS / [MS + IBS]), which is the portion of the photosynthetic part of the leaf that can be responsible for refixing carbon obtained through the C_4_ cycle. Finally, we also measured the distance between bundle sheaths (BSD) and width of the inner bundle sheath (IBSW) using the mean widths of equatorial cells.

**Figure 1:**
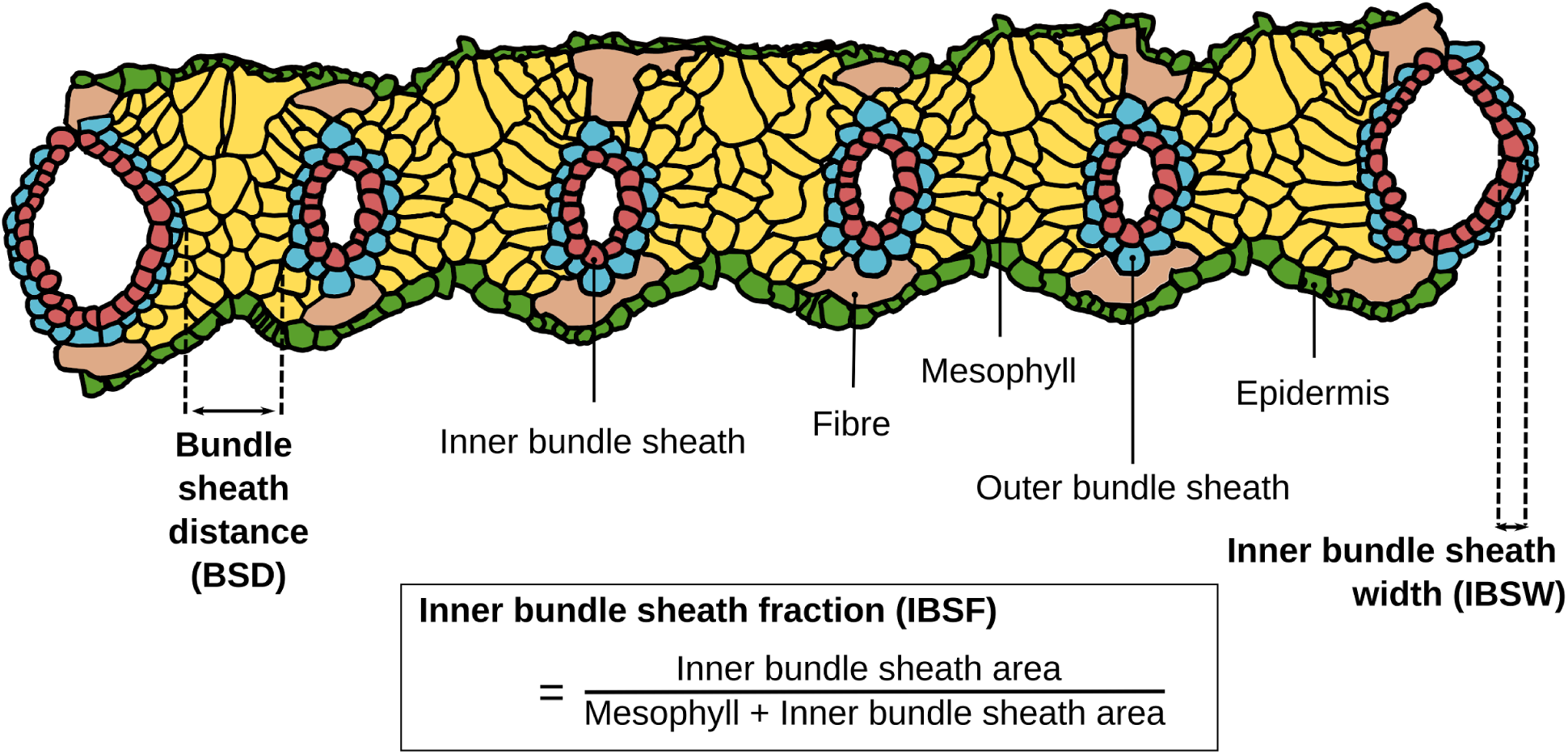
A cross-section of a typical C_4_ *Alloteropsis semialata* leaf. The schematic is traced from a cross-section of accession JKO23-03_16, with tissue types labeled. The anatomical measurements used for the genome-wide association study (GWAS) are indicated in bold.

### Estimating trait heritability

To estimate the proportion of phenotypic variation explained by underlying genetic differences, we calculated the heritability of δ^13^C (complete and restricted C_3_+C_4_ datasets) and the leaf anatomical traits (C_3_+C_4_ dataset) using Genome-wide Complex Trait Analysis (GCTA) v.1.94.1 (Yang et al., 2011). A genetic relationship matrix was inferred from the previously generated SNP calls and combined with the phenotype values in GCTA. Heritability was then estimated for each trait using the restricted maximum likelihood (REML) method.

### Genome wide association study

We performed a genome-wide association study (GWAS) for several photosynthetic traits, with the objective of ultimately proposing some candidate genes underpinning the phenotype. We used the variation in photosynthetic type which exists across *A. semialata* as a whole, before focusing on anatomical variation in the C_3_+C_4_ accessions that has been associated with the strength of the C_4_ cycle (Alenazi et al., 2023). We defined our associated regions of the genome as the linkage block containing a significant SNP from the GWAS. We then identified the gene models located within the correlated region as candidate genes, and assessed their functional relevance, gene expression patterns and selective forces they have been evolving under.

The GWAS itself was performed using the rMVP package (Yin et al., 2021) in R studio v.4.3, with the MVP.Data function and default parameters used for single-locus GWAS analysis for each phenotypic trait with the fixed and random model circulating probability unification (FarmCPU) approach (Yin et al. 2021). Population structure and genetic relatedness can confound a GWAS and result in false associations (Chen et al. 2016). We therefore included the previously generated pairwise kinship matrix so that the relationships among samples could be accounted for. The phenotypic data for each trait was normalised (if required) and a Bonferroni corrected SNP significance threshold of p ≤ 0.05 was used.

### Linkage disequilibrium

Linkage blocks are regions of the genome that are likely to be co-inherited, and the association of the significant SNPs identified from the GWAS could be caused by any gene within this region. To determine the linkage block encapsulating each SNP we used Haploview v.4.1 (Barrett et al., 2005). The input map and binary files were processed using PLINK-v1.9 (Purcell et al. 2007), and we used a solid spine of LD with default parameters to infer linkage block size (Kim et al. 2018). This approach requires the first and last SNPs in a block to be in strong LD with all intermediate markers (normalised deviation [D’] ≥ 0.8), but the intermediate markers do not necessarily need to be in LD with each other. Identifying linkage blocks is heavily impacted by the distribution of SNPs across the genome, something that is accentuated by reduced sequencing methods such as ddRADSeq. We therefore used the genome-wide mean linkage block size if the analysis failed to place a significant SNP in a block of its own. To do this, we positioned the significant SNP at the center of the artificial linkage block, and if necessary truncated it to avoid incorporating unlinked SNPs up and/or downstream from this marker.

### Identification of candidate genes

The linkage blocks associated with the phenotype of interest contain the causal gene(s) in addition to those that happen to be in close physical linkage (hitchhiking). To try and identify plausible candidate genes in each region we compared their functional annotations, expression patterns and the selective pressures they are evolving under.

Orthofinder v.2.5.4 (Emms and Kelly, 2015) was used to identify orthologous genes to the loci in the associated regions. To do this, we combined the *A. semialata* protein sequences with nine other plant species (*Arabidopsis thaliana*, *Brachypodium distachyon*, *Hordeum vulgare*, *Oryza sativa*, *Physcomitrium patens*, *Solanum lycopersicum*, *Triticum aestivum,* and *Zea mays*) downloaded from Phytozome v.13 (Goodstein et al., 2012). We then used publicly available databases (e.g. TAIR [Berardini et al., 2000], RAP-DB [Sakai et al., 2013], and maizeGDB [Monaco et al., 2013]) and literature searches to extrapolate the functions of each orthogroup containing a gene from a correlated linkage block identified from the GWAS.

Gene expression data for the candidate genes was extracted from a phylogenetically informed gene expression study of C_4_ evolution in *A. semialata* (Dunning et al., 2019a) to determine the expression pattern of the candidate genes. We also used these data to test for differential expression between the photosynthetic types using two-tailed t-tests, with p-values Bonferroni corrected to account for multiple testing.

Finally, we used whole-genome resequencing data (Bianconi et al., 2020) for 45 *A. semialata* accessions to determine if the genes in the GWAS regions were evolving under positive selection. In short, the datasets were downloaded from NCBI sequence read archive, mapped to the reference genome using bowtie2 and consensus sequences were generated using previously developed methods (Olofsson et al., 2016; Dunning et al., 2022) and a maximum-likelihood phylogeny tree for each gene was inferred using RAxML (Stamatakis, A., 2014) with 100 bootstrap. We then inferred the selective pressure each gene was evolving under by running the M0 model in codeML v.4.9h (i.e. a single *dN/dS* ratio for all branches and sites).

## Results

### Population structure

The broadscale phylogenetic (Figure 2a) and population genetic (Figure 2b) analyses recovered those previously inferred by earlier studies, with the different photosynthetic types (C_3_, C_3_+C_4_ and C_4_) belonging to separate clades (Olofsson et al., 2016, 2021; Bianconi et al., 2020). Within the C_3_+C_4_ intermediates, accessions are separated into five populations geographically spread across the Central Zambezian miombo woodlands (Figure 2). This reconfirms the phylogenetic groupings previously demarcated (Alenazi et al., 2023), although the earliest diverging sixth lineage is absent in this study because it is only represented by a single herbarium accession from the Democratic Republic of the Congo and lacks ddRADSeq data. The population structure analysis (Figure 2c) largely concurs with the phylogenetic groupings, although it indicates gene-flow between populations. The distribution of the C_3_+C_4_ groups has a pattern largely matching a scenario of isolation-by-distance along an east-west axis through Zambia and Tanzania (Figure 2d).

**Figure 2:**
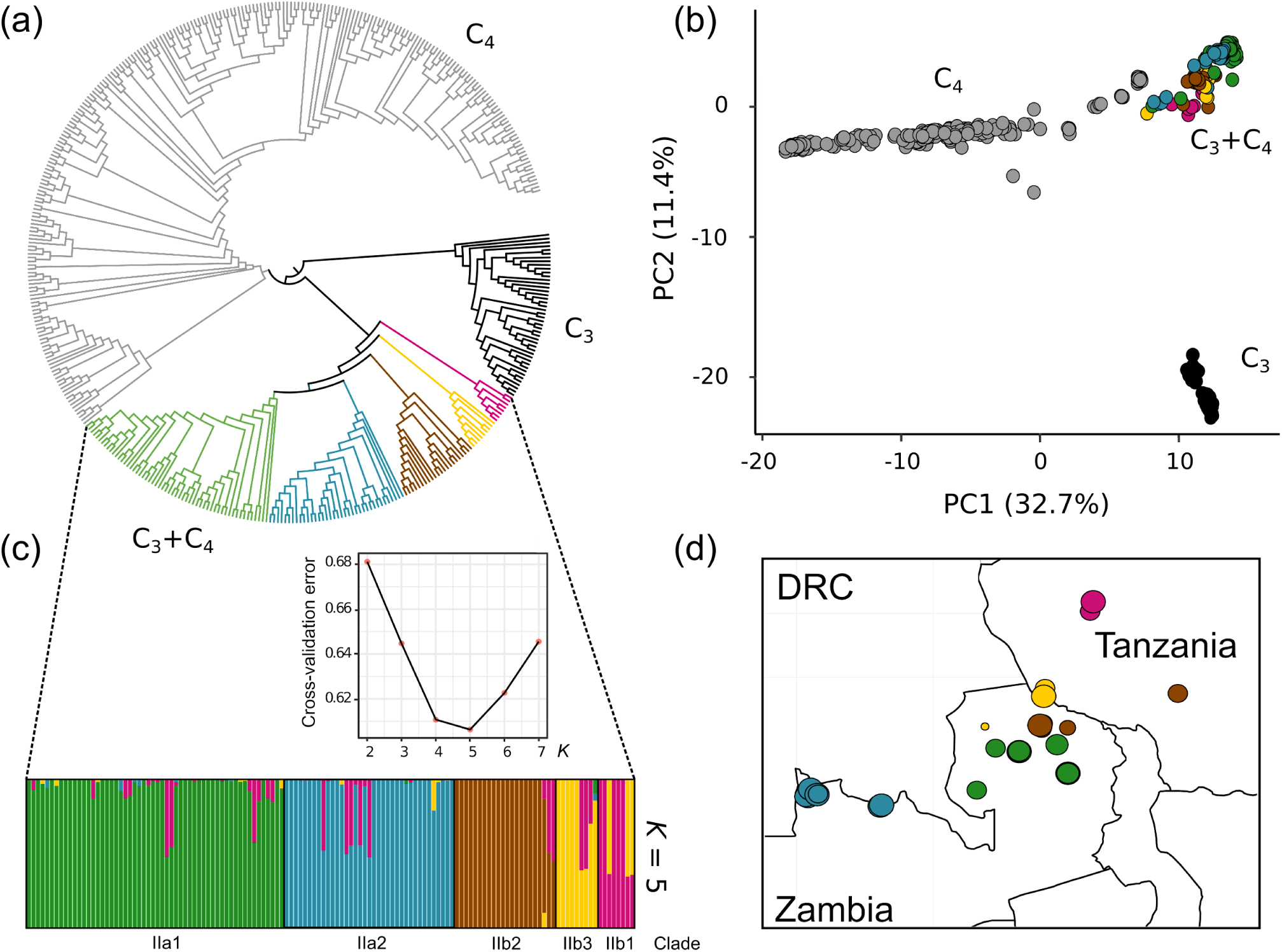
Population genomics of the *Alloteropsis semialata* accessions. (a) A cladogram of the maximum likelihood phylogenetic tree, with individual clades recovered within the C_3_+C_4_ lineage coloured (same colours used in all panels). (b) A principal component analysis of the genotypes, showing the first two axes. (c) Admixture results for the C_3_+C_4_ *A. semialata* accessions for *K* = 5, the optimal number of population clusters based on the cross-validation error. (d) location of the C_3_+C_4_ populations used in this study, with the size of the point proportional to the number of samples (range 1-20 samples per population).

### Identifying regions of the genome correlated with the strength of the C_4_ cycle

We used the δ^13^C values as a proxy for the strength of the C_4_ cycle for all 420 *A. semialata* samples used in this study. As expected, the δ^13^C values supported the demarcation of the main nuclear clades into the C_3_, C_3_+C_4_ and C_4_ phenotypes (Figure 3a). For C_3_ and C_4_ accessions, we found δ^13^C average values of -26.67 and -12.63 with little dispersion within each group, whereas for C_3_+C_4_ accessions, we found substantial variation ranging from -28.35 to -18.47 with an average of -23.87. The heritability estimate, which represents the proportion of phenotypic variation due to genetic variation in the population, was high for δ^13^C when considering all photosynthetic types (h^2^ = 0.75; SE = 0.06; n = 420), and three-fold lower when just considering the C_3_+C_4_ intermediates (h^2^ = 0.25; SE = 0.00; n = 132).

**Figure 3.**
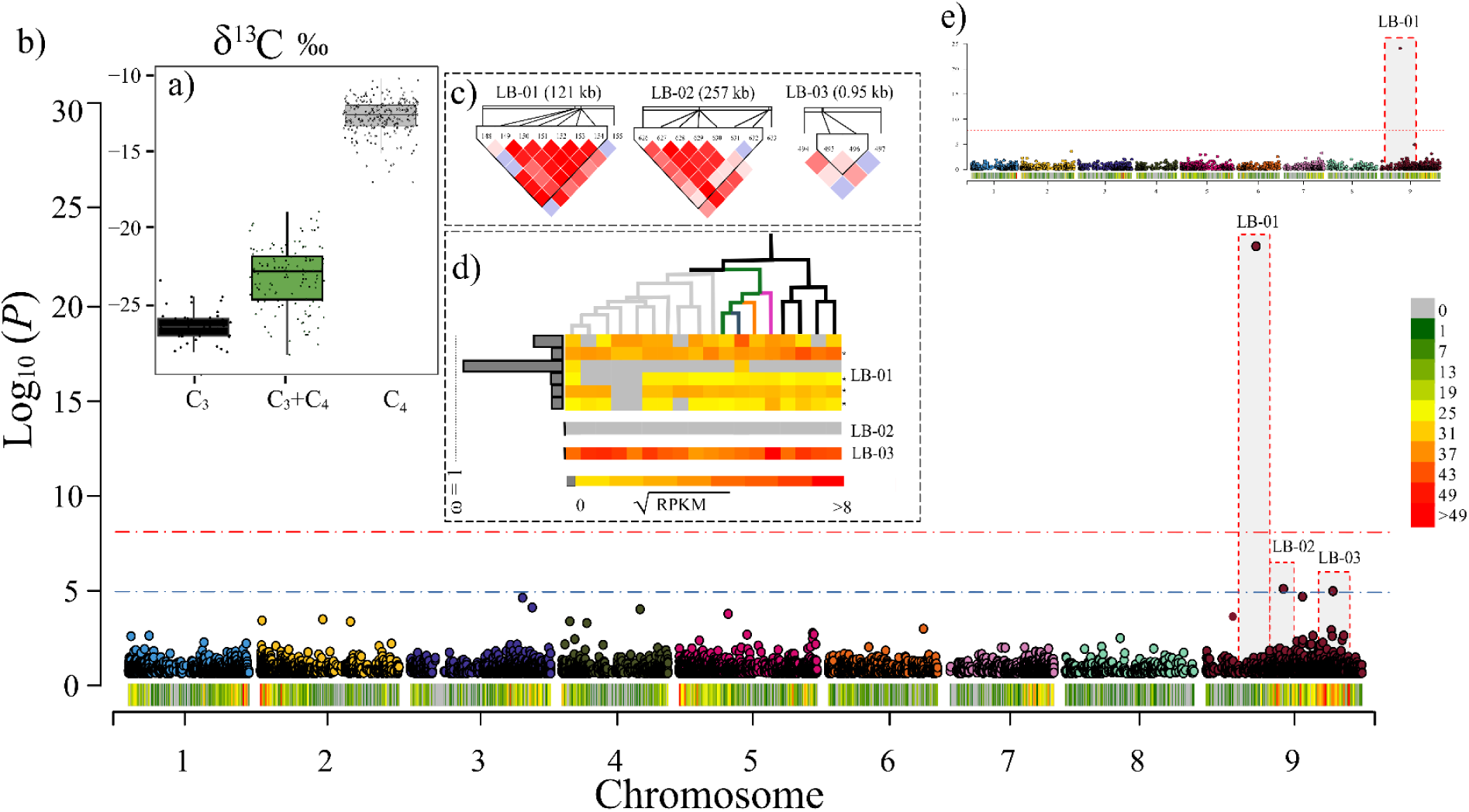
Genetic variation associated with the strength of the C_4_ cycle in *Alloteropsis semialata*. (a) The stable carbon isotope ratio (δ^13^C) was used to infer the strength of the C_4_ cycle, with values measured for each of the photosynthetic types shown. The boxes show the median value and the interquartile range, and the whiskers represent 1.5× the interquartile range. (b) Manhattan plot showing the results of a Genome-Wide Association Study (GWAS) for δ^13^C using all samples. The blue and red dotted lines indicate Bonferroni corrected P-values of 0.05 and 0.001, respectively. Significant SNPs are labeled with a block ID. The density of markers is shown along each chromosome on the x-axis. (c) Heatmap of pairwise linkage disequilibrium (LD) between markers surrounding each significant SNP, ranging from white indicating low LD (LOD < 2 and D’ < 1) to bright red indicating strong LD (LOD ≥ 2 and D’ = 1). (d) For each of the genes in each linkage block we show their expression level and selective pressure they are evolving under. The heat map shows square-root transformed leaf expression levels extracted from Dunning et al., (2019), ordered based on the phylogenetic relationships (grey = C_4_, black = C_3_, other = individual C_3_+C_4_ clades). The bars at the side of the heat map indicate the omega value for each gene (grey bar ω < 1; red bar ω > 1). (e) shows the results of the GWAS analysis when only using the C_3_+C_4_ accessions.

We conducted a combined GWAS using all samples (Figure 3b), as well as various partitions by photosynthetic type (Figure S2). When considering all accessions, the GWAS identified three significant SNPs on chromosome 9, which all corresponded to relatively narrow regions based on the LD (Figure 3c). The region with the highest association with δ^13^C (LB-01) is a 121 kb region at 32.2 Mb (Table 1, Table S3 and Figure 3b). The same region was also significant when repeating the GWAS within the C_3_+C_4_, and when combining the C_3_+C_4_ with either the C_3_ or C_4_ accessions (Table S3, Figure 3b and Figure S2), but not when excluding the C_3_+C_4_ accessions. These results imply that the underlying causative gene segregates only within the C_3_+C_4_ group. There were six predicted protein coding genes in the LB-01 region, and all were expressed in the leaf tissue of at least one *A. semialata* accession (Table S4). One of these genes (SLP*1* [ASEM_AUS1_34305]) was significantly more highly expressed in the C_3_ than in the other photosynthetic types (C_3_ vs C_4_ Bonferroni adjusted P-value = 0.073; C_3_ vs C_3_+C_4_ Bonferroni adjusted P-value = 0.015; Table S4), although there is no consistent differential expression between photosynthetic types when individual populations are compared separately (Dunning et al. 2019a). None of the six genes were found to be strictly evolving under positive selection with a dN/dS ratio (ω) > 1 (Figure 3d), although those with the highest values may be seeing a relaxation of purifying selection (e.g. ω = 0.83 for ASEM_AUS1_34303). The annotated genes in the LB-01 region have a variety of functions (Table S4), including loci associated with the regulation of the Calvin cycle (*SLP1* [ASEM_AUS1_34302]) and the activation of NADP-malic enzyme 2 (NADP-ME2), a C_4_ decarboxylation enzyme (RIPK [ASEM_AUS1_34305]).

**Table 1:** Significantly correlated regions of the genome identified in the genome wide association studies.

| Phenotype | Chromosome | SNP position | LD block (kb) | $-\log_{10} P$ | Bonferroni adjusted $P$-value | Number of genes |
| --- | --- | --- | --- | --- | --- | --- |
| $\delta^{13}\text{C}$ | 9 | 32191256 | 121 | 29.18 | 5.33E-26 | 6 |
|  | 9 | 49592498 | 0.95** | 5.73 | 1.51E-02 | 1 |
|  | 9 | 81051792 | 63 | 5.58 | 2.12E-02 | 1 |
| IBSF | 9 | 58663539 | 362 | 6.88 | 3.58E-04 | 33 |
|  | 2 | 634463 | 6** | 4.83 | 4.95E-02 | 2 |
|  | 5 | 23291361 | 59* | 5.89 | 3.53E-03 | 4 |
|  | 4 | 63231217 | 13 | 6.71 | 5.34E-04 | 2 |
|  | 8 | 23357684 | 275 | 16.47 | 9.25E-14 | 24 |
| BSD | 9 | 10612628 | 11 | 5.1 | 2.17E-02 | 1 |
|  | 9 | 48749591 | 0.14** | 5.59 | 7.02E-03 | 0 |

\* This SNP was not located in a linkage block in our analyses, we therefore defined the region using the genome wide median block size
\*\* This SNP was not located in a linkage block in our analyses, we therefore defined the region using the genome wide median block size that was truncated if there was a closely located unlinked SNP up or downstream.

The two other regions identified in the δ^13^C GWAS using all samples (LB-02 and LB-03; Figure 3b) were not significant when partitioning the data by photosynthetic type (Supplementary Table S3). Both these regions are delimited by LD blocks narrow in size and that contain one annotated gene each. The candidate gene in LB-02 was not expressed at all in any *A. semialata* mature leaves, while the one in LB-03 was expressed in all accessions, but was not differentially expressed between photosynthetic types. In addition, both genes do not seem to have been under positive selection (Figure 3d). One of these genes (ASEM_AUS1_29467) encodes a SCARECROW-LIKE protein 9 (SCL9) protein belonging to the GRAS gene family, a group of transcription factors shown to play a key role in C_4_ leaf anatomy and photosynthetic development in maize (Slewinski et al., 2012; Hughes & Langdale, 2020). The other gene encodes a protein associated with the suppression of non-photochemical quenching and maintaining the efficiency of light harvesting (*SOQ1* [ASEM_AUS1_14480]).

### Identifying regions of the genome associated with C_4_ leaf anatomy in the C_3_+C_4_ intermediates

We studied the genetic basis of three leaf anatomical traits previously associated with the strength of the C_4_ cycle (δ^13^C) using the132 C_3_+C_4_ individuals (Figure 1; Alenazi et al., 2023). The heritability estimates for the three leaf anatomical traits in the C_3_+C_4_ intermediates ranged from roughly equivalent to the value for δ^13^C value to much lower (IBSF h^2^ = 0.22 [SE = 0.04]; BSD h^2^ = 0.12 [SE = 0.06]; IBSW h^2^ = 0.06 [SE = 0.06]; n = 132). No significantly correlated genomic region was detected for inner bundle sheath width (IBSW) (Figure S3), the trait with the lowest heritability. However, we did detect SNPs significantly associated with bundle sheath distance (BSD) and inner bundle sheath fraction (IBSF).

#### i. Bundle sheath distance (BSD)

The distance between consecutive bundle sheaths (BDS) plays a significant role in determining the rate and efficiency of photosynthesis in plants, with smaller distances being significantly correlated with higher δ^13^C (more C_4_-like) values (Alenazi et al., 2023). The C_3_+C_4_ intermediate accessions showed a range of BSDs from 55.14 to 178.36 μm, with variation between subclades (Figure 4a). The GWAS identified two significant regions associated with BSD, both on chromosome 9 (Table 1; Table S3; Figure 4). Only one annotated gene was identified in the correlated genomic regions associated with BSD, the function of which is associated with leaf development (*GSL8* [ASEM_AUS1_16831]; Table S4).

**Figure 4.**
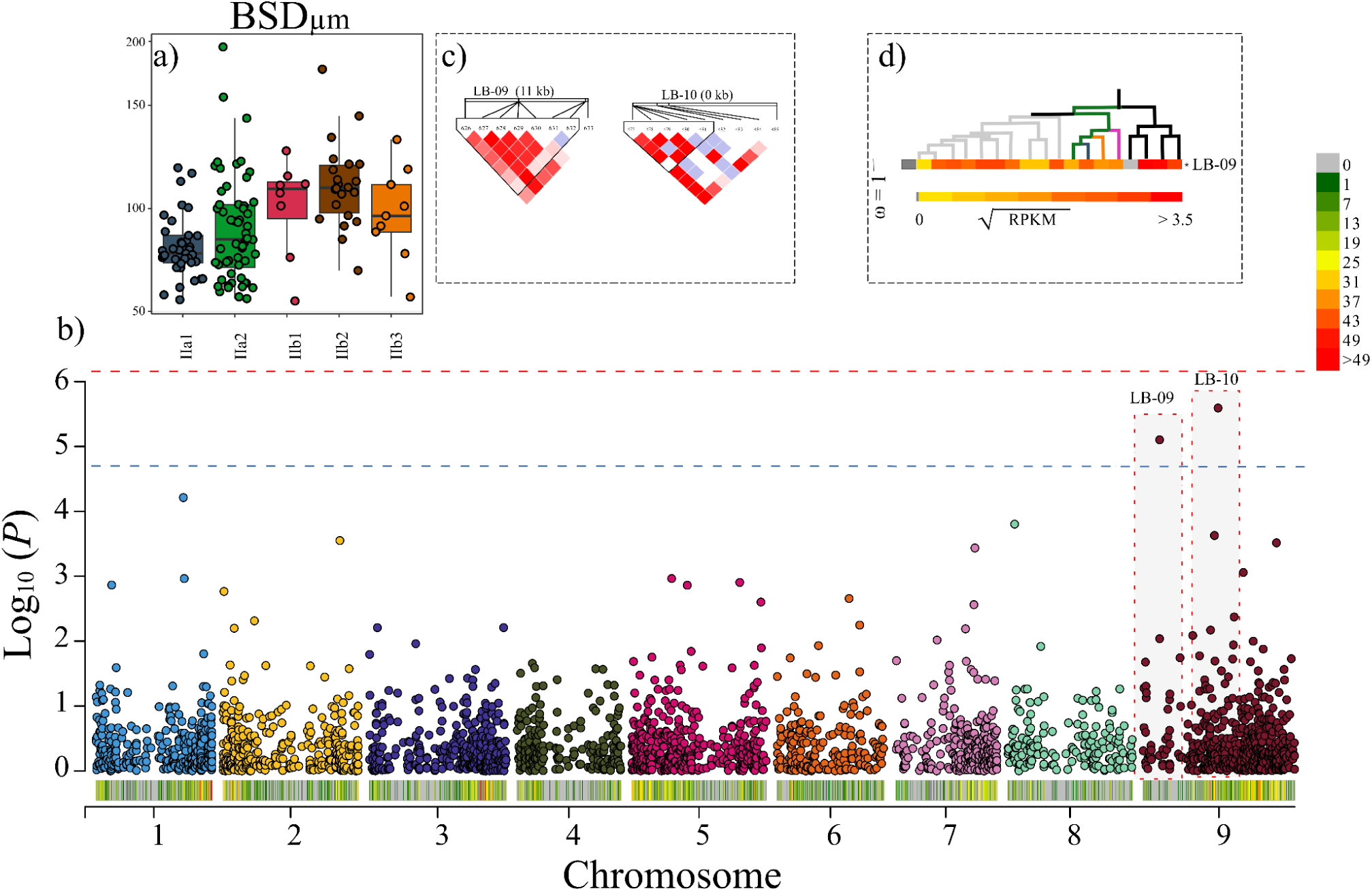
Genetic variation associated with bundle sheath distance (BSD) in the C_3_+C_4_ *Alloteropsis semialata*. (a) The boxplot shows the BSD variation for each of the C_3_+C_4_ subclades. The box indicates the median value and the interquartile range, and the whiskers represent 1.5× the interquartile range. (b) A Manhattan plot showing the results of a Genome-Wide Association Study (GWAS) for BSD. The blue and red dotted lines indicate Bonferroni corrected P-values of 0.05 and 0.001, respectively. Significant SNPs are labeled with a block ID. The density of markers is shown along each chromosome on the x-axis. (c) Heatmap of pairwise linkage disequilibrium (LD) between markers surrounding each significant SNP, ranging from white indicating low LD (LOD < 2 and D’ < 1) to bright red indicating strong LD (LOD ≥ 2 and D’ = 1). (d) For each of the genes in each linkage block we show their expression level and selective pressure they are evolving under. The heat map shows square-root transformed leaf expression levels extracted from Dunning et al., (2019), ordered based on the phylogenetic relationships (grey = C_4_, black = C_3_, other = individual C_3_+C_4_ clades). The bars at the side of the heat map indicate the omega value for each gene (grey bar ω < 1; red bar ω > 1).

#### ii. Inner bundle sheath fraction (IBSF)

Inner bundle sheath fraction (IBSF) represents the portion of the leaf that can be used for C_4_ photosynthesis (Figure 1). A higher IBSF in the C_3_+C_4_ *A. semialata* has been significantly correlated with a higher δ^13^C (more C_4_ like) (Alenazi et al., 2023). In the C_3_+C_4_ populations, there is a range from 0.05 to 0.29, with variation between subclades (Figure 5a). We identified five regions of the genome correlated with IBSF, each on a different chromosome (Table 1 & S3, and Figure 5). Expression was detected in mature leaves for 62% of the 65 genes located in the five regions, with no consistent differential expression between photosynthetic types in mature leaves (Dunning et al. 2019a), although two are on average more highly expressed in the C_3_ vs C_4_ accessions (*YlbH* [ASEM_AUS1_21119] Bonferroni adjusted P-value = 0.073; *STR12* [ASEM_AUS1_17094] Bonferroni adjusted P-value = <0.001). 5 out of the 65 genes were also evolving under strong positive selection with a dN/dS ratio (ω) > 1 using the one-ratio model (Table S4). The annotated genes in the correlated regions of the genome have a variety of functions (Table 1, Table S4 and Figure 6), including loci directly connected to the response to light stress (*FAH1* [ASEM_AUS1_36251]) and leaf development (*GAT19* [ASEM_AUS1_21136], *CNOT11* [ASEM_AUS1_25789] & *GRF1* [ASEM_AUS1_21151]).

**Figure 5.**
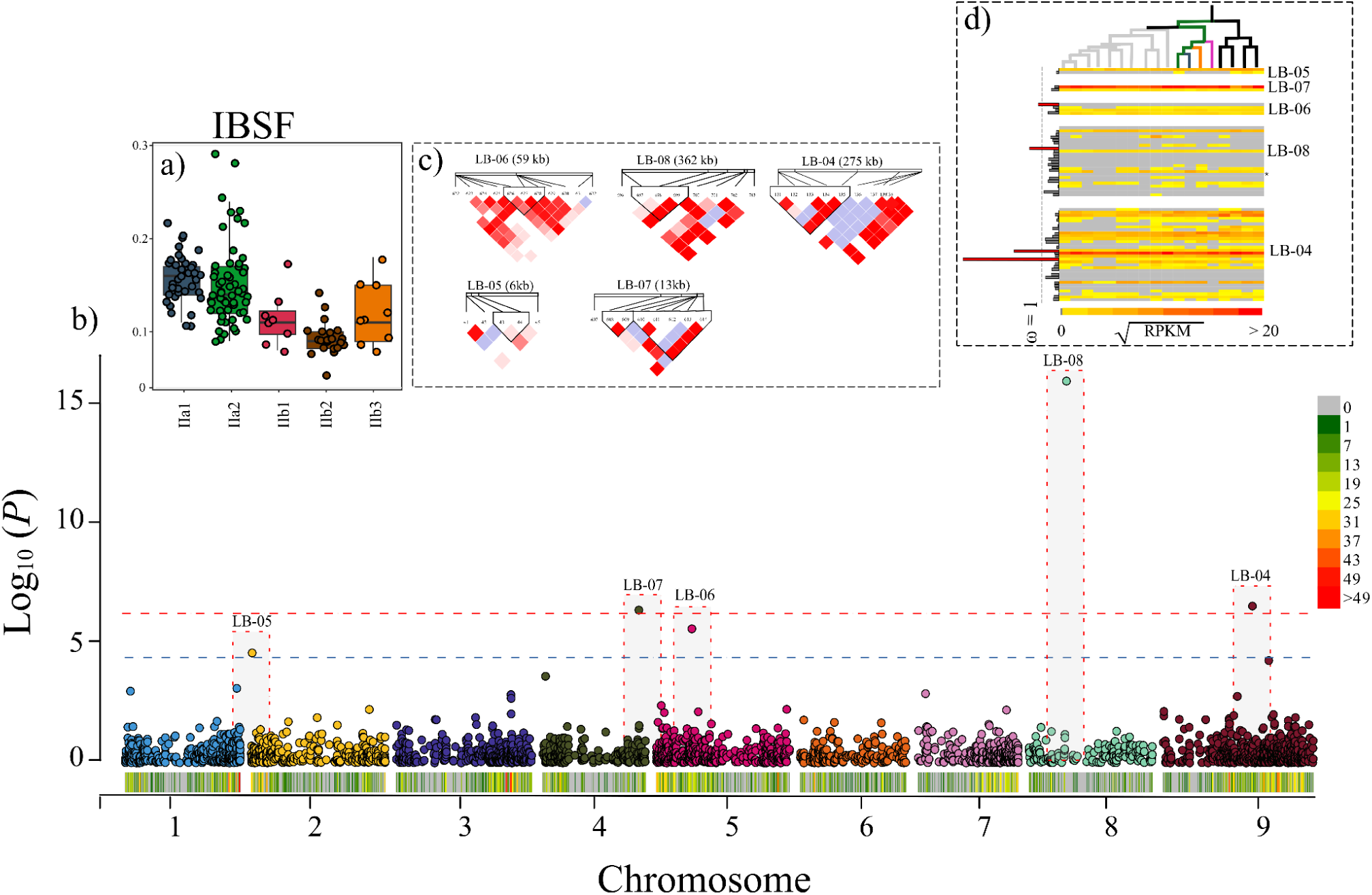
Genetic variation associated with inner bundle sheath fraction (IBSF) in the C_3_+C_4_ *Alloteropsis semialata*. (a) The boxplot shows the IBSF variation for each of the C_3_+C_4_ subclades. The box indicates the median value and the interquartile range, and the whiskers represent 1.5× the interquartile range. (b) a Manhattan plot showing the results of a Genome-Wide Association Study (GWAS) for IBSF. The blue and red dotted lines indicate Bonferroni corrected P-values of 0.05 and 0.001, respectively. Significant SNPs are labeled with a block ID. The density of markers is shown along each chromosome on the x-axis. (c) Heatmap of pairwise linkage disequilibrium (LD) between markers surrounding each significant SNP, ranging from white indicating low LD (LOD < 2 and D’ < 1) to bright red indicating strong LD (LOD ≥ 2 and D’ = 1). (d) For each of the genes in each linkage block we show their expression level and selective pressure they are evolving under. The heat map shows square-root transformed leaf expression levels extracted from Dunning et al., (2019), ordered based on the phylogenetic relationships (grey = C_4_, black = C_3_, other = individual C_3_+C_4_ clades). The bars at the side of the heat map indicate the omega value for each gene (grey bar ω < 1; red bar ω > 1).

## Discussion

C_4_ photosynthesis is a remarkable example of convergent evolution that has facilitated certain plants to adapt to high temperatures. *Alloteropsis semialata* is the only known species with C_3_, C_3_+C_4_ and C_4_ genotypes. It is therefore a useful model to study the initial steps leading to the establishment of the C_4_ phenotype since these modifications are not conflated with other changes that accumulate over time (Pereira et al., 2023), and its emergence in this species provided an immediate demographic advantage (Sotelo et al., 2024). Here, we estimate the heritability and identify regions of the genome correlated with variation in both the stable carbon isotope ratio (δ^13^C) and leaf anatomical traits known to influence δ^13^C (Alenazi et al., 2023). Despite a relatively modest sample size (n = 420 for δ^13^C; n = 132 for leaf anatomy), we identified regions of the genome significantly associated with these traits, which indicate that the genetic architecture of C_4_ evolution in *A. semialata* is relatively simple.

### Genetic basis of the carbon isotope ratio (δ^13^C) in Alloteropsis semialata

Using linked phenotype and genotype information for 420 *A. semialata* individuals, we identified three associated regions of the genome, containing seven protein coding genes (Figure 3). The underlying differences in the δ^13^C between photosynthetic types is driven by C_4_ plants evolving to fix carbon with the PEPC enzyme rather than Rubisco. However, genes encoding PEPC were not detected in the associated regions identified in our GWAS. This absence could be due to variation in the specific PEPC gene copy used for C_4_ in the individual accessions masking the signal, with up to five different versions known to be used by different *A. semialata* accessions (Dunning et al., 2017). Among these five copies, three were laterally acquired (Christin et al., 2012), complicating the matter further as they appear as large structural variants inserted randomly into the genome (Dunning et al., 2019b), and are only present in a subset of individuals (Raimondeau et al., 2023). However, based on the annotations of the genes in the associated regions, we did identify candidate genes with functions potentially associated with the δ^13^C, the most promising of which include those associated with the regulation of Rubisco (*SLP1* [ASEM_AUS1_34302]), the activation of the NADP-ME C_4_ decarboxylating enzyme (*RIPK* [ASEM_AUS1_34305]), the development of C_4_ ‘Kranz’ anatomy (*SCL9* [ASEM_AUS1_29467]), and the suppression of non-photochemical quenching (*SOQ1* [ASEM_AUS1_14480]).

*SLP1* encodes a Shewanella-like protein phosphatase 1, an ancient chloroplast phosphatase (Johnson et al., 2020) that is generally more highly expressed in photosynthetic tissue (Kutuzov and Andreeva, 2012). In *Arabidopsis thaliana*, it is co-expressed with a number of photosynthetic genes (including all of the Calvin cycle enzymes and Rubisco activase) and it is predicted to play a role in the light-dependent regulation of chloroplast function (Kutuzov and Andreeva, 2012). In *A. semialata*, *SLP1* is significantly more highly expressed in the C_3_ accessions compared to the other photosynthetic types. This greater expression in C_3_ accessions could indicate a higher Calvin cycle activity at the whole leaf level, meanwhile in the C_3_+C_4_ and C_4_ individuals its expression would be increasingly restricted to the inner bundle leaf tissue. Subdivision of the light signaling networks is one of the key steps in the partitioning of photosynthesis across tissue types in C_4_ species (Hendron & Kelly, 2020), and *SLP1* is potentially one of the regulators of this key innovation in *A. semialata*.

RIPK is an enzyme that plays a role in disease resistance and plant immunity (Liu et al., 2011), but has pleiotropic effects. In *A. thaliana,* RIPK directly phosphorylates NADP-ME2 to enhance its activity and increase cytosolic NADPH concentrations (Wu et al., 2022). In C_4_ species, CO_2_ is initially fixed in the mesophyll by CA and PEPC before being transported to an internal leaf compartment and released for Rubisco to assimilate through the Calvin cycle. Preliminary studies in *A. semialata* concluded that NADP-ME was the predominant decarboxylating enzyme, although its activity varied with temperature (Frean et al., 1983). Subsequent transcriptome work showed that NADP-ME expression has a mean expression level four times higher in C_4_ and C_3_+C_4_ accessions (mean = 300 RPKM; SD = 235) than in C_3_ plants (mean = 75 RPKM; SD = 32), although this difference is not always consistent between populations (Dunning et al., 2019a). The other decarboxylating enzyme commonly used by C_4_ *Alloteropsis* accessions is phosphoenolpyruvate carboxykinase (PCK), but like PEPC, a C_4_ copy of PCK was also laterally acquired (Christin et al., 2012), complicating its identification in a GWAS analysis because it is absent in the C_3_ accessions (Dunning et al., 2019b).

*SCL9* belongs to the GRAS gene family of transcription factors that regulate plant development (Hirsch & Oldroyd, 2009). This multigene family includes two known C_4_ Kranz anatomy regulators identified in maize, SHORTROOT (Slewinski et al., 2014) and SCARECROW (Slewinski et al., 2012). Orthologous SCARECROW (*SCR*) genes have divergent functions, being recruited for distinct roles in leaf development within maize, rice and *A. thaliana* (Hughes & Langdale, 2022). In addition to its influence on leaf anatomy, SCR is also required for maintaining photosynthetic capacity in maize (Hughes & Langdale, 2020). The correlation of the SCARECROW-LIKE *SCL9* gene with the strength of the C_4_ cycle in *A. semialata* may indicate that convergence in C_4_ phenotypes are a result of the parallel recruitment of GRAS transcription factors between species, although there is divergence in the specific loci recruited for this purpose.

*SOQ1* is a chloroplast-localized thylakoid membrane protein that regulates non-photochemical quenching in *A. thaliana* (Duan et al., 2023). In full sunlight, plants absorb more light energy than they can process, which can ultimately result in the generation of free radicals that damage the photosynthetic apparatus (Müller et al., 2001). To overcome this, plants have evolved non-photochemical quenching which enables them to dissipate the excess energy as heat. This problem is potentially exacerbated in C_4_ species, which typically grow in high-light conditions compared to their C_3_ counterparts (Sage and Monson, 1999). Preliminary evidence indicates that C_4_ species exhibit a significantly faster and greater non-photochemical quenching relaxation than their C_3_ relatives, including between photosynthetic types in *A. semilata* (Arce Cubas, 2023). *SOQ1* may therefore play a direct role in regulating differences in the non-photochemical quenching responses among *A. semialata* photosynthetic types, and it may represent a good candidate gene to target for reduced photoinhibition associated with fluctuating light conditions in crops (Long et al., 1994)

### The genetic basis of C_4_ leaf anatomy

In *A. semialata,* the inner bundle sheath is the site of C_4_ photosynthesis, and three leaf anatomical variables linked to the proliferation of this tissue explain the strength of the C_4_ cycle (δ^13^C) in the C_3_+C_4_ intermediate accessions: inner bundle sheath width (IBSW), bundle sheath distance (BSD) and inner bundle sheath fraction (IBSF) (Alenazi et al., 2023). IBSW has the lowest heritability (h^2^ = 0.06 [SE = 0.06]), and we failed to identify any significant SNPs correlated with this phenotype in our GWAS. This absence of significant genetic factors contributing to the trait may indicate that IBSW has a complex genetic architecture or high phenotypic plasticity. The δ^13^C can be influenced by environmental effects on water-use efficiency, and the previously observed IBSW correlation with δ^13^C may potentially arise from such environmental induced plasticity (Alenazi et al., 2023). For example, bundle sheath cells in wheat have a larger diameter (more C_4_-like) under drought conditions (Osonubi et al., 2017).

### Plasmodesmata and reduced distance between bundle sheaths

We identified two regions of the genome associated with BSD that contain a single protein coding gene. This gene is *GSL8* (ASEM_AUS1_16831), a member of the Glucan Synthase-Like (GSL) family that encodes enzymes synthesising callose. *GSL8* plays an important role in tissue-level organisation (Chen et al., 2009), including stomatal (Guseman et al., 2010) and leaf vein patterning (Linh and Scarpella, 2022). Mutants of *GSL8* in *A. thaliana* formed networks of fewer veins in their leaves (Linh and Scarpella, 2022). This change in venation is mediated by the aperture of plasmodesmata, channels through cell walls that connect neighbouring cells (Paterlini, 2020; Band, 2021), which is regulated by GSL8 (Saatian et al., 2018; Linh and Scarpella, 2022). Normal vein patterning is reliant on an auxin hormone signal traveling through these plasmodesmata, and any interference of this signal disrupts leaf vein development (Linh and Scarpella, 2022). GSL8 might play a role in strengthening the C_4_ cycle in *A. semialata* by reducing the distance between bundle sheaths through modulation of the auxin signal. The transition to being fully C_4_ in *A. semialata* is also correlated with the presence of minor veins, which reduces both the number of mesophyll cells and the distance between bundle sheaths in C_3_+C_4_ in comparison with C_3_ populations (Lundgren et al., 2019). Therefore, GSL8 may play a pleiotropic role in the strengthening of C_4_ photosynthesis in *A. semialata* by increasing both the proportion of bundle sheath tissue in the leaf, and the connectivity between the two distinct cell types required to complete the cycle.

### The genetic basis of the inner bundle sheath fraction in Alloteropsis semialata

The inner bundle sheath fraction (IBSF) has the highest heritability of all the three leaf anatomy measures used (h^2^ = 0.22 [SE = 0.04]). Since it is a composite trait, it is more likely to be influenced by multiple developmental processes. Our GWAS identified five regions of the genome significantly associated with IBSF, containing 65 predicted protein coding genes. Interestingly, we found a number of genes associated with leaf development that could play a role in the development of C_4_ leaf architecture. These include homologs of genes that alter leaf area and vascular development (*GATA transcription factor 19* [ASEM_AUS1_21136]) (An et al., 2020), leaf thickness (*CCR4-NOT transcription complex subunit 11* [ASEM_AUS1_25789]) (Sarowar et al., 2007), and leaf width by regulating meristem determinacy (GRF1-interacting factor 1 [ASEM_AUS1_21151]) (Zhang et al., 2018). *GRF1* (also called *ANGUSTIFOLIA3*) is perhaps the most interesting of these genes, since it is expressed in the mesophyll cells of leaf primordium and can influence the proliferation of other clonally independent leaf cells (e.g. epidermal cells [Kawade et al., 2013]). The numerous regulators of leaf development identified in the GWAS point to an interacting balance of growth regulators to increase the proportion of bundle sheath tissue within the leaf for C_4_ photosynthesis.

There are other genes in these regions with a diverse set of functions, although it is unclear how they could modulate IBSF, including genes associated with light stress and lignin biosynthesis. *FAH1* encodes ferulic acid 5-hydroxylase (F5H) 1, a cytochrome P450 protein that, when disrupted, reduces anthocyanin accumulation under photooxidative stress (Maruta et al., 2014) and is more highly expressed in the C_3_ (mean RPKM = 7.38; SD = 5.29) than other photosynthetic types (mean RPKM = 1.00; SD = 2.10; Table S4). These loci could also play a role in C_4_ photosynthesis, although most likely they might just be in close physical linkage.

## Conclusion

C_4_ photosynthesis is a complex trait that requires the rewiring of metabolic gene networks and alterations to the internal leaf anatomy. Identifying the genetic basis of these key innovations can be complicated by the divergence time between C_3_ and C_4_ comparisons. Here, we exploited the photosynthetic diversity within *Alloteropsis semialata*, the only known species to contain C_3_, C_3_+C_4_ intermediate, and C_4_ phenotypes, to identify the genes underlying this transition. We first performed a GWAS analysis for the strength of the C_4_ cycle using δ^13^C as a phenotype, and identified regulators of C_4_ decarboxylation enzymes (*RIPK*), non-photochemical quenching (*SOQ1*), and photosynthetic development (*SCARECROW-LIKE*). We then conducted a GWAS for leaf morphological traits linked to the δ^13^C in the C_3_+C_4_ intermediates and identified several genes involved in tissue-level organisation and leaf development that could underpin the proliferation of C_4_ bundle sheath tissue in *A. semialata*. Overall, the detection of genomic regions significantly associated with C_4_ traits with a relatively modest sample size points to a relatively simple genetic basis of the C_4_ syndrome in *A. semilata*, and the candidate genes highlighted here represent ideal loci to investigate with follow-up functional studies.

## Data availability

All *A. semialata* genomic data was previously published, and the additional phenotype data generated here is available in the SI.

## Author contributions

ASA, LP, PAC, CPO and LTD designed the study. ASA conducted the experimental work and generated the phenotype data. ASA, LP & LTD analysed the data. All authors interpreted the results and helped write the manuscript.

## Acknowledgments

ASA is supported by a PhD scholarship from the Northern Border University in Saudi Arabia, LP is supported by a Natural Environment Research Council grant NE/V000012/1, PAC was funded by a Royal Society University Research Fellowship (grant URF\R\180022), and LTD is funded by a NERC fellowship (grant NE/T011025/1).

