## Supplementary material for "Identifying leaf anatomy and metabolic regulators that underpin C_4_ photosynthesis in *Alloteropsis semialata*": SI

**Supplementary Figures:** 3

**Supplementary Tables:** 4

**Supplementary Datasets:** 2

### Supplementary Figures

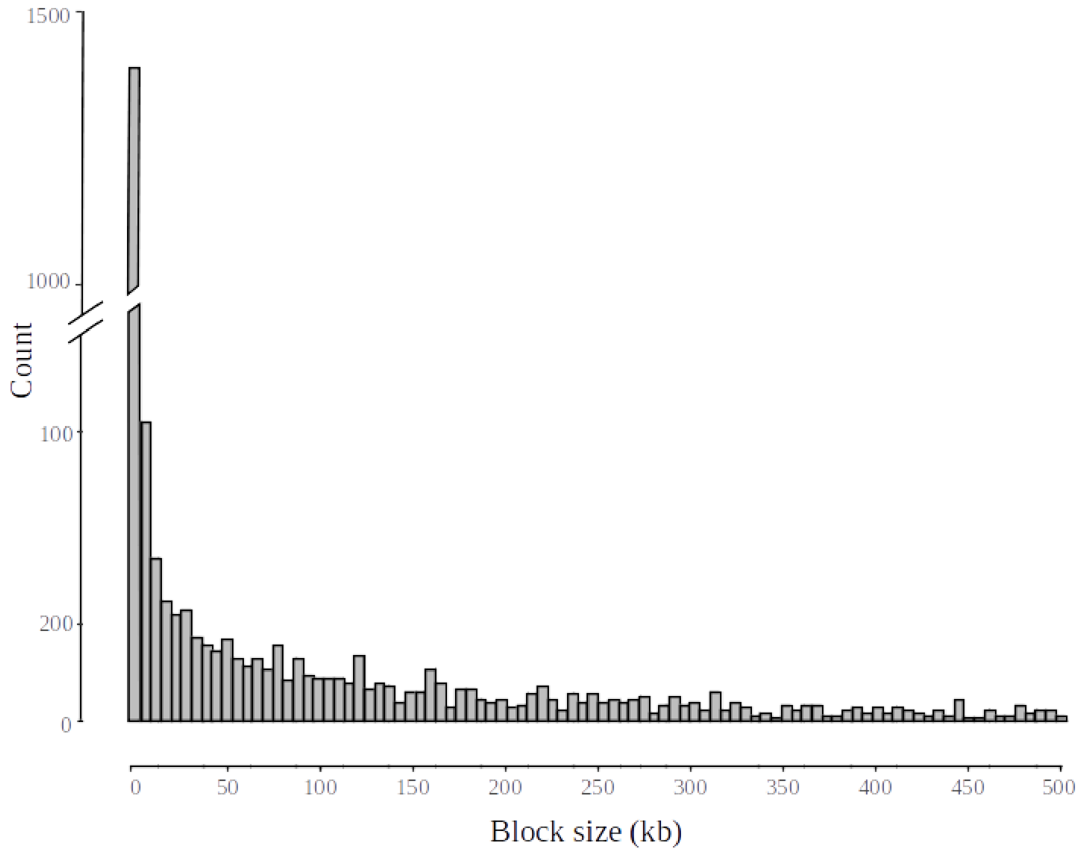


**Figure S1: Distribution of linkage block sizes in the *Alloteropsis semialata* genome.** The block sizes were inferred using a solid spine of linkage disequilibrium (SSLD) analysis. 2,601 linkage blocks were identified, with a mean length of 58,996 kb (SD = 109,378 kb).

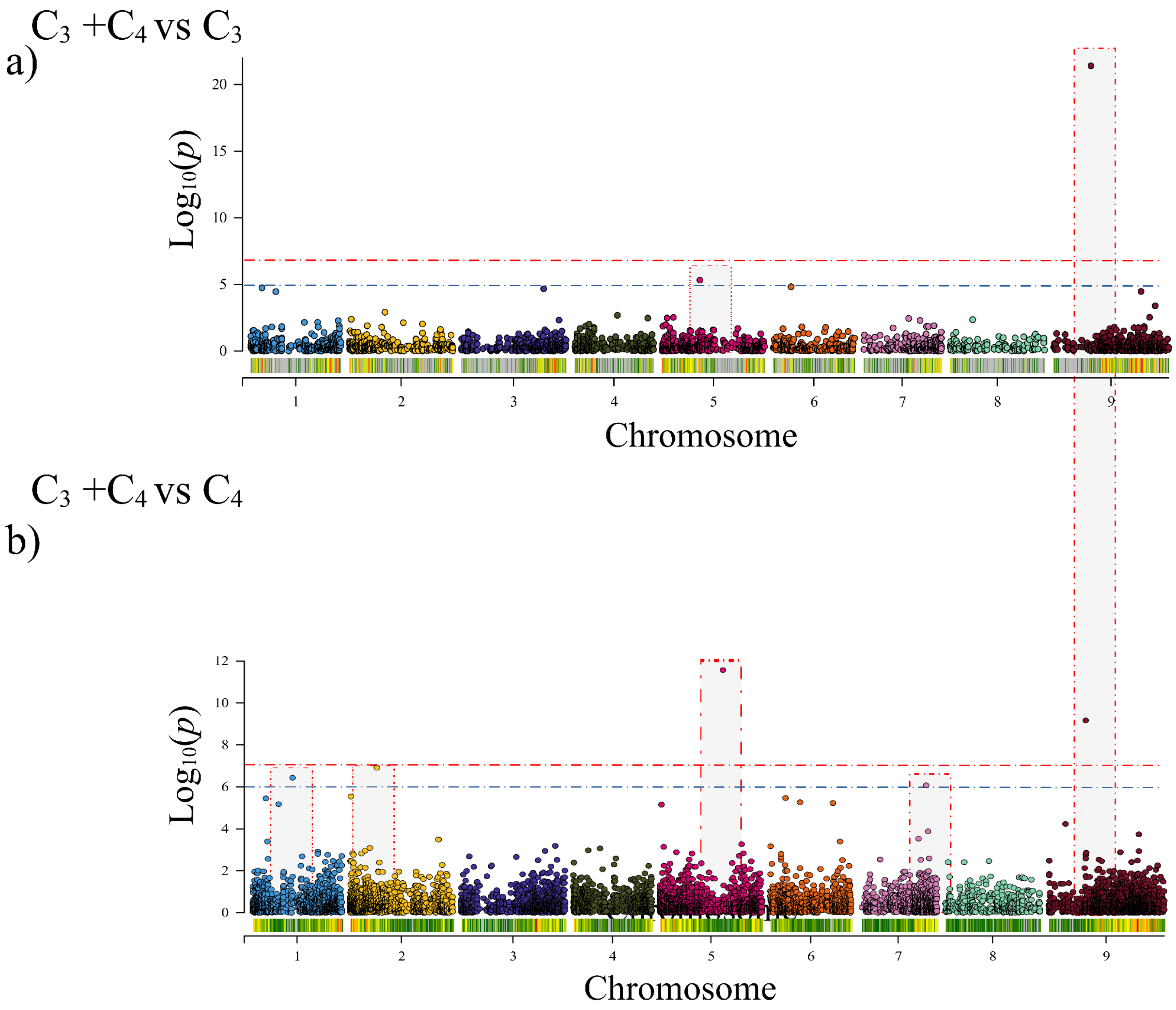
a)
C3 +C4 vs C3
Log10(p)
0
5
10
15
20
1
2
3
4
5
6
7
8
9
Chromosome
b)
C3 +C4 vs C4
Log10(p)
0
2
4
6
8
10
12
1
2
3
4
5
6
7
8
9
Chromosome

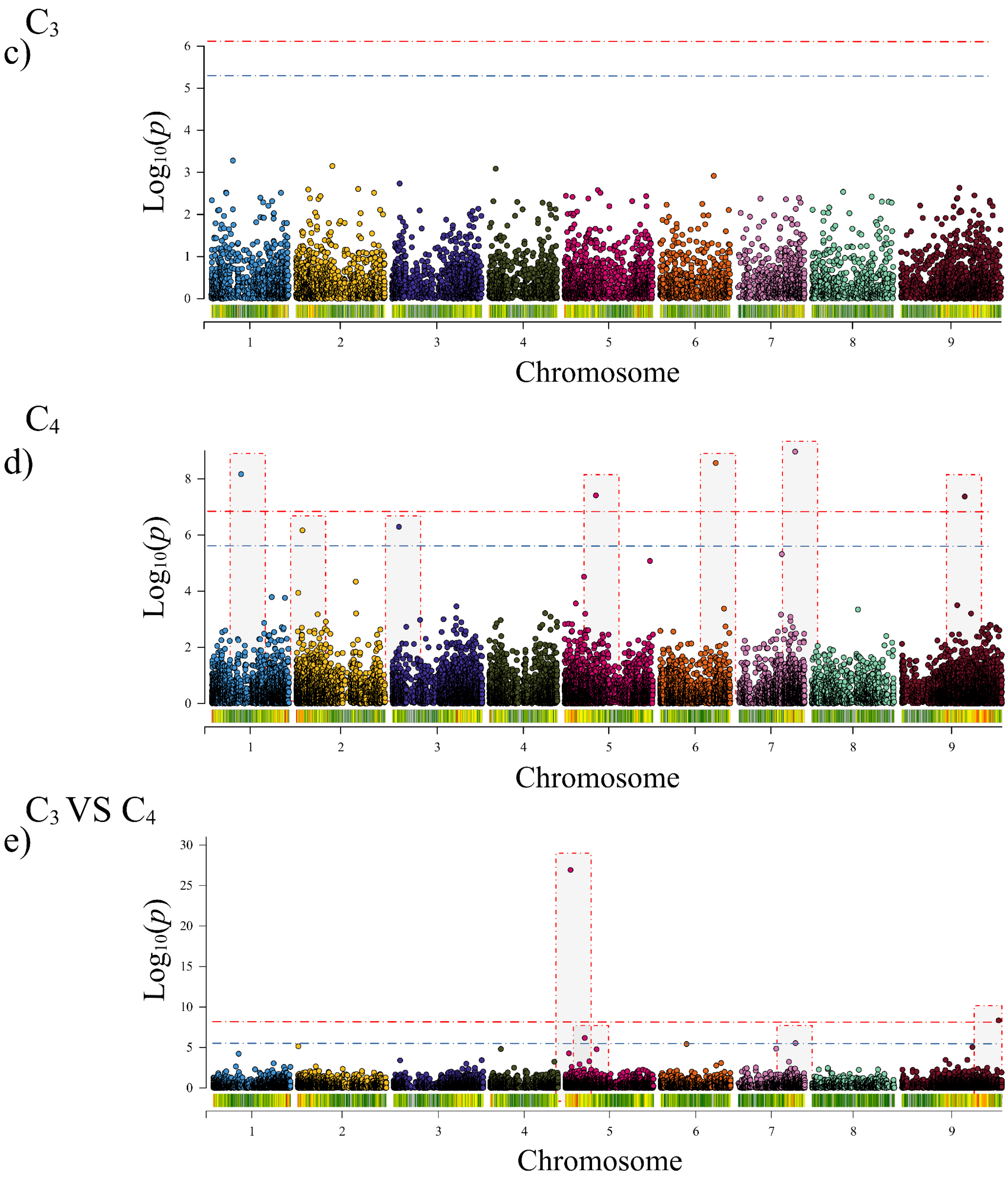


**Figure S2: Manhattan plot showing the results of a Genome-Wide Association Study (GWAS) for δ13C using various subdivisions of samples based on photosynthetic type.**

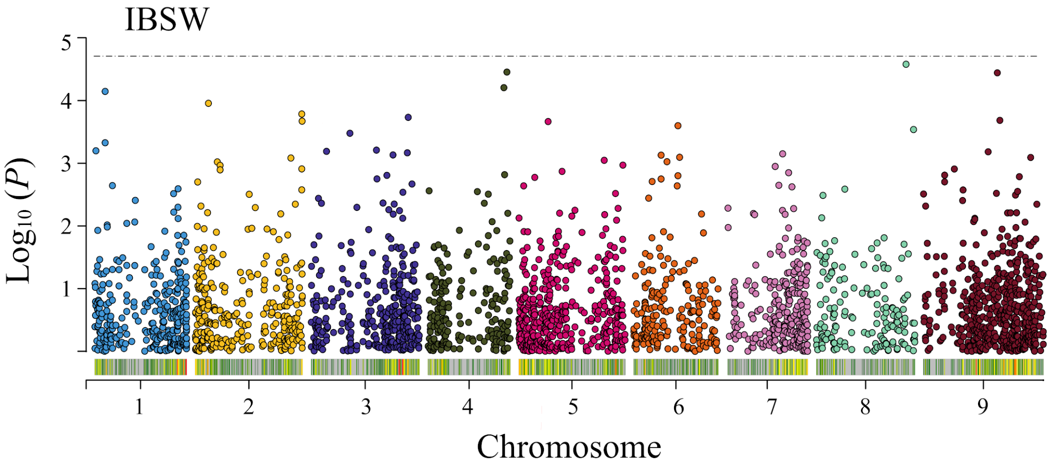


**Figure S3: Manhattan plot showing the results of a Genome-Wide Association Study (GWAS) for inner bundle sheath width (IBSW).** The blue dotted line indicates a cutoff significance value of a Bonferroni corrected P-value of 0.05.

Supplementary Information

### Supplementary Tables

*All tables available as individual tabs in separate excel file*

**Table S1: Details of samples used in the genome wide association study.** This includes sample collection information, $\delta_{13}C$ and NCBI Sequence Read Archive accession numbers.

**Table S2: Leaf anatomy measurements for the $C_3$+$C_4$ *Alloteropsis semialata*.**

**Table S3: Summary of significant regions detected.** This includes the SNP location and linkage block size.

**Table S4: Summary of genes in significant regions.** This includes gene annotation, expression levels and estimates of omega (dN/dS ratio).

### Supplementary datasets

**Data set 1: Maximum likelihood phylogeny of all accessions used in this study.**

https://drive.google.com/file/d/1L6mkIvT7JSXMcOuixxTILqnnB-v-J5p4/view?usp=drive_link

**Data set 2: Orthogroup phylogenies**

https://drive.google.com/file/d/1KgHxTg3PXNAQIkvW3lQRtokivDKuGAAX/view?usp=drive_link
